# A distribution-aware and functionally relevant novel framework for generation and discovery of bioactive peptides

**DOI:** 10.64898/2026.08.04.742799

**Authors:** Rachit Abhigyan, Vikas Sood, Pooja Arora, Baljeet Kaur

**Affiliations:** Department of Computer Science, Hansraj College, University of Delhi; Department of Biochemistry, Jamia Hamdard, Delhi; Department of Zoology, Hansraj College, University of Delhi

**Keywords:** Bioactive peptides, Peptide generation, Generative-Evolutionary framework, Transformer, Evolutionary optimization, Class imbalance

## Abstract

Recent advances in artificial intelligence have accelerated the discovery of bioactive peptides by enabling computational exploration of the vast peptide sequence space. However, existing peptide generation approaches generally rely on either distribution-learning models, which generate biologically realistic sequences but do not consistently optimize functional activity, or optimization-based methods, which maximize prediction confidence while often deviating from the underlying distribution of experimentally validated peptides. To address this limitation, a two-phase generative–evolutionary framework is proposed that integrates distribution learning with evolutionary optimization. In the first phase, Variational Autoencoders (VAE), Autoregressive Transformers (ART), and Token Diffusion Transformers (TDT) are used to generate biologically plausible seed peptides. In the second phase, these peptides were used as initial seed for Hill Climbing optimization procedure that iteratively improves fitness function score. The proposed two-phase framework was evaluated using a dataset of experimentally validated IL-2-inducing peptides. Evaluation using independent IL-2 prediction models showed that Autoregressive Transformer combined with Hill Climbing achieved the best overall performance, achieving the mean IL-2 induction confidence score of 0.96 while reducing KL divergence from 2.26 for standalone Hill Climbing to 0.75. A case study on an independent IL-13 inducing peptide dataset showed similar trends, with ART initialized Hill Climbing achieving the mean IL-13 induction score of 0.99 while reducing KL divergence from 1.76 to 0.59. Overall, the framework provides a generalizable approach for balancing functional optimization and distributional realism and can be applied to peptide discovery and data augmentation in imbalanced biological datasets thereby generating high confidence peptides for wet lab validation.

**Highlights:**

- Proposed a two-phase framework for bioactive peptide generation with potential to address class imbalance in peptide classification tasks.
- Performed a systematic comparison of distribution-learning and optimization-based approaches for peptide generation.
- Combined distribution-learning models for sequence generation with optimization algorithms for improving peptide functional properties.
- Demonstrated the applicability of the proposed framework across multiple bioactive peptide datasets.

## 1. INTRODUCTION

The discovery of bioactive peptides has emerged as a powerful and promising approach for the development of next-generation therapeutics, including antimicrobial, antiviral, and immunomodulatory agents [1]. Due to their high specificity, favorable safety profiles, and relatively low toxicity, peptides have emerged as promising alternatives to conventional small-molecule drugs. More than 100 peptide-based therapeutics have been approved for clinical use worldwide, with many others currently undergoing clinical evaluation [2]. As a result, peptide-based drug discovery has attracted increasing research interest. Despite these advances, the discovery of novel bioactive peptides remains challenging. The peptide sequence space is extremely large, making exhaustive experimental screening infeasible. For example, a peptide of length 20 has 20^20^ possible sequences, making comprehensive enumeration and evaluation impractical. In addition, the limited availability of experimentally validated peptide datasets restricts the development of computational models for peptide prediction and design. Recent advances in artificial intelligence, particularly deep generative models, have provided new approaches for peptide generation. Distribution-learning models, including Variational Autoencoders (VAEs), Autoregressive Transformers, and diffusion-based models, have been used to learn peptide sequence distributions and generate novel peptide sequences [3–5]. Trained on experimentally validated bioactive peptides, these models generate realistic and diverse sequences while exploring new regions of the peptide sequence space. A recent comparative study [6] showed that no single generative approach consistently outperformed the others, with different models exhibiting strengths in distributional similarity, sequence diversity, and sequence exploration. However, these models focus on learning the overall peptide sequence distribution and do not explicitly optimize generated sequences for specific functional properties. Therefore, distribution-learning approaches alone may not be sufficient for targeted peptide design. To overcome this limitation, optimization-based methods inspired by evolutionary principles, such as Hill Climbing and Genetic Algorithms [7], have been applied to directly optimize peptide sequences for predefined objectives. By introducing mutations and selecting high-performing variants, these methods explore the sequence space and identify peptides with improved functional scores. However, optimization-based methods also have certain limitations. They are often biased towards local optima and may not adequately capture the broader sequence distribution, resulting in reduced diversity and possible overfitting to specific sequence motifs. Moreover, without an understanding of the global peptide distribution, optimization methods may generate sequences that have high functional scores but still deviate from biologically plausible patterns. An effective peptide design would benefit by integrating the distribution learning and functional optimization in a single framework. To bridge this gap, this work proposes a novel two-phase framework to leverage the ability of distribution-learning models, to learn realistic sequence distributions, with the capacity of optimization algorithms to enhance functional properties. This work has exhaustively explored well known distribution-learning algorithms, namely, Variational Autoencoders, Autoregressive Transformers, and Token Diffusion Transformers along with the popular and effective evolutionary based optimization algorithms, viz, Hill algorithm and Genetic Algorithm to generate novel peptide sequences with high distribution fidelity and biological activity.

The findings demonstrate distinct and complementary support of these two approaches in the proposed two-phase framework. The distribution-learning models, VAE and the two Transformer-based architectures, when implemented as standalone models, though successfully reproduced the sequence distribution, they did not achieve comparable functional optimization. Also, it was observed that optimization-based methods, Hill Climbing and Genetic Algorithm, when implemented individually, were effective in increasing functional scores but failed to accurately capture the underlying distribution of bioactive peptides. Importantly, the proposed two-phase framework combines these strengths, generating peptide sequences that are both high-scoring and distribution-consistent. The proposed framework was evaluated on the dataset from recently published studies aiming to classify IL-2 [8, 9] inducing peptides. We observed that our two-phase approach consistently outperformed standalone optimization-based and standalone distribution-learning approaches in both prediction performance and distributional alignment of IL-2 inducing peptides.

Further, a case study was conducted on the IL-13-inducing peptide dataset reported in previous studies [10, 11] to evaluate the generalizability of the proposed framework. The observed trends were consistent with those obtained for the IL-2 dataset, demonstrating that the proposed framework is robust across datasets of different sizes.

In addition to peptide generation, the proposed framework has potential applications in addressing class imbalance in peptide prediction tasks. Machine learning models are often trained on datasets in which experimentally validated bioactive peptides represent a relatively small positive class [11–13]. This imbalance can bias the learning process towards the majority class, reducing the ability of models to identify rare bioactive peptides. The problem is particularly relevant in peptide prediction tasks, where experimentally validated positive samples are often limited.

Several data-level and algorithm-level approaches have been proposed to address class imbalance, including random oversampling [14], Adaptive Synthetic Sampling (ADASYN) [15], random undersampling [16], and synthetic sample generation methods such as SMOTE [17]. However, these approaches are not directly applicable to biological sequence data because they do not account for the sequence and physicochemical constraints of peptides. As a result, they may generate unrealistic or biologically implausible sequences that do not reflect the properties of experimentally validated peptides [18]. The proposed framework provides a biologically informed alternative by generating synthetic peptides that preserve the characteristics of real bioactive peptides while increasing the size and diversity of the positive class.

Overall, this study presents a two-phase framework that combines distribution learning with evolutionary optimization for bioactive peptide generation. The framework generates peptide candidates with improved predicted functional properties while preserving the characteristics of experimentally validated peptides. Although demonstrated on IL-2–inducing peptides and validated on an independent IL-13 dataset, the framework is general and can be applied to other peptide classes, including antimicrobial, antiviral, and immunomodulatory peptides.

## 2. MATERIAL AND METHODS

### 2.1 Dataset

The dataset used in this study was obtained from the experimentally validated IL-2–inducing and non-inducing peptides reported in the recently published IL2PepScan [9]. The original dataset consisted of 4,411 IL-2–inducing peptides (positive samples) and 4,076 non-inducing peptides (negative samples). Standard preprocessing was performed to improve data quality and consistency. Peptides containing non-standard amino acids (B, J, O, U, X, and Z) were removed. Duplicate sequences within each class were eliminated, and peptides present in both the positive and negative sets were removed from both classes. A length distribution analysis was performed, and only peptides between 8 and 30 amino acids were retained. After preprocessing, the final dataset contained 3,385 IL-2–inducing peptides and 3,050 non-inducing peptides. Only the experimentally validated IL-2–inducing peptides were used to train the generative models so that they could learn the distribution of bioactive peptide sequences. The non-inducing peptides were retained for reference and evaluation where required.

### 2.2 Feature Extraction

The machine learning models used in this study require numerical input features. Therefore, the peptide sequences were converted into numerical representations using two widely used and high-performing feature extraction methods.

#### 2.2.1 Dipeptide Composition (DPC)

Dipeptide Composition (DPC) [19] represents a peptide sequence based on the normalized frequency of all possible adjacent amino acid pairs. For a sequence of length *L*, there are *L* − 1 overlapping dipeptides, resulting in a 400-dimensional feature vector (20 × 20 possible pairs).

The DPC feature for a dipeptide (*i*, *j*), is defined as:

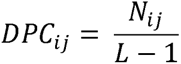

where *N_ij_* is the number of occurrences of dipeptide (*i*, *j*), in the sequence, and *L* is the sequence length. DPC captures local compositional patterns and was used as the feature representation for the evaluation prediction model.

#### 2.2.2 Dipeptide Deviation from Expected Mean (DDE)

Dipeptide Deviation from Expected (DDE) [19] measures the deviation of observed dipeptide frequencies from their expected values based on amino acid composition and codon usage.

For a dipeptide (*i*, *j*), the expected frequency is defined as:

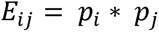

where *p_i_* and *p_j_* represent the probabilities of occurrence of amino acids *i* and *j*, respectively. The probability of each amino acid *p_i_* was computed based on codon usage, where 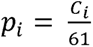, and *C_i_* represents the number of synonymous codons encoding amino acid *i*.

The DDE feature is computed as:

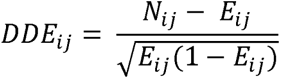

where *N_ij_* is the observed frequency of dipeptide (*i*, *j*).

DDE captures deviations from expected compositional patterns and was used as the feature representation for defining the fitness function in optimization-based methods.

### 2.3 Distribution-Learning Models

#### 2.3.1 Variational Autoencoder

A Variational Autoencoder (VAE) [20, 21] was implemented to learn latent space representation and model the distribution of IL2-inducing peptides. The model was trained on experimentally labelled positives that were recently used in IL2PepScan study [9]. The sequences were padded to a length of 30 amino acids and represented using one-hot encoding over a vocabulary consisting of 20 natural amino acids and a padding token. The encoder is composed of a LSTM layer with 128 hidden units, followed by two parallel fully connected layers that output the 16-dimensional mean and log-variance vectors of a multivariate gaussian latent distribution. Latent vectors were sampled using the reparameterization trick to enable gradient-based optimization. The decoder consisted of a Repeat Vector layer followed by an LSTM with 128 units and a Time Distributed Dense layer with Softmax activation to reconstruct amino acid probabilities at each position. The loss function used was a sum of categorical cross-entropy reconstruction loss and Kullback-Leibler (KL) divergence regularization, encouraging latent representations to approximate standard normal distribution. Temperature sampling was applied to modulate diversity during sequence generation. Unique peptides with lengths between 8 and 30 were collected until 2000 sequences were obtained.

#### 2.3.2 Autoregressive Transformer

An Autoregressive Transformer [22, 23] (ART) was implemented to learn conditional probability distribution of IL2-inducing peptides. It learns 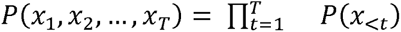, that is, each amino acid is predicted based on all previous amino acids. Hence, the generation process works autoregressively. The model was trained on experimentally validated positive sequences. Each peptide was prepended with a start token and padded to a fixed length of 30 amino acids. Sequences were represented as integer-encoded tokens. The model architecture was composed of a 64-dimensional embedding layer, followed by learned positional embeddings for encoding sequence order. Two stacked transformer blocks were employed, each consisting of multi-head self-attention (4 heads) with causal masking followed by a feed-forward network with 128 hidden units. Residual connections, layer normalization and dropout were applied to stabilize training. The model was trained using categorical cross-entropy loss to maximize the likelihood of predicting the next amino acid conditioned on preceding tokens. This approach explicitly captures sequential dependencies. Generation begins with start token and proceeds token by token until padding token is produced or maximum length is achieved. Temperature sampling was applied to modulate diversity during decoding.

#### 2.3.3 Token Diffusion Transformer

A discrete diffusion-based transformer model [24] (TDT) was implemented to generate IL2-inducing peptides via iterative denoising i.e., reverse diffusion process. The padding and integer-encoding was utilized as before over a vocabulary consisting of 20 natural amino acids, a padding token, and a mask token. During training, a forward corruption process was defined by masking tokens within valid sequence positions. At diffusion timestep t, each amino acid was replaced with probability t/T independently where T = 20 represents the total number of diffusion steps. Padding positions were excluded from corruption. The denoising model was conditioned on both the corrupted sequence and timestep. Self-attention was applied without causal masking, allowing each position to attend to the full corrupted sequence during denoising. The architecture consisted of token embeddings, learned positional embeddings, and learned timestep embeddings, each 64-dimensional which were summed to form the input representation. Two stacked transformer blocks, each comprising multi-head self-attention with 4 heads and feed-forward layers of 128 hidden units, were used. Residual connections and layer normalization were used to stabilize training. The model was trained to predict the original clean sequence from its corrupted version by minimizing sparse categorical cross-entropy loss computed only over masked positions. This objective encourages the model to learn the reverse diffusion process, that is, iterative reconstruction of clean sequences from progressively corrupted inputs. For generation, sequences were initialized as fully masked tokens with pre-defined lengths sampled from training length distribution. Reverse diffusion was performed iteratively from timestep T to 1. At each step, a subset of masked positions was probabilistically updated based on the model’s predicted token distributions, while padding tokens were preserved. After completing all diffusion steps, remaining mask tokens were removed to obtain the final peptide sequences. Unique peptides within the specified length range were collected until 2000 sequences were generated.

### 2.4 Evolutionary Optimization Methods

#### 2.4.1 Hill Climbing

Hill climbing (HC) is a classical local search-based optimization algorithm [25] which optimizes a robust fitness function that involves IL-2 confidence score. It is utilized for generation of sequences by generating a randomized variable length sequence sampled uniformly from 20 natural amino acids, mutating it 200 times and updating sequence if a mutation increases the fitness function score, that is, IL-2-induction confidence score as predicted using IL2PepScan model [9]. The mutation consists of random amino acid substitution, insertion or deletion. A strictly greedy acceptance criterion was used i.e. a mutation was accepted only if it strictly increases the confidence score. Duplicate generation is handled.

#### 2.4.2 Genetic Algorithm

Genetic Algorithm (GA) is a population based evolutionary optimization-centric approach [26]. It has been widely applied in peptide and protein design tasks [27]. In this study, it was implemented with an initial population of 200 randomly generated peptides. A fitness function was used which is defined as the IL-2 confidence score. A global archive was maintained to store unique peptides encountered during evolution. Elitism was applied by directly inheriting the top 10% of the population into next generation. The remaining population was generated using crossover and mutation. Parent sequences were generated randomly from top 50 ranked sequence thereby favoring high-fitness candidates. Offspring were generated using single-segment crossover where prefix on one parent was appended with suffix of another with a mutation probability of 0.2. The new population replaced previous generation except for the elite sequences. Evolution proceeded iteratively until 2000 sequences were collected in global archive or for a maximum of 300 generations.

### 2.5 Proposed Two-Phase Framework

The proposed framework integrates distribution learning with evolutionary optimization through a two-phase pipeline for bioactive peptide generation. It first generates biologically realistic seed peptides using a distribution-learning model and then refines these sequences using an evolutionary optimization algorithm guided by a task-specific fitness function. This allows the framework to combine the strengths of both approaches by generating peptides that follow the underlying sequence distribution while improving their functional properties. Although the framework is compatible with a wide range of distribution-learning and optimization algorithms, this study uses Variational Autoencoders (VAE), Autoregressive Transformers (ART), and Token Diffusion Transformers (TDT) as representative distribution-learning models. Among the optimization algorithms evaluated, Hill Climbing demonstrated the best empirical performance and was therefore selected as the optimization component of the proposed framework. Figure 1 provides a schematic representation of the proposed two-phase framework.

**Figure 1.**
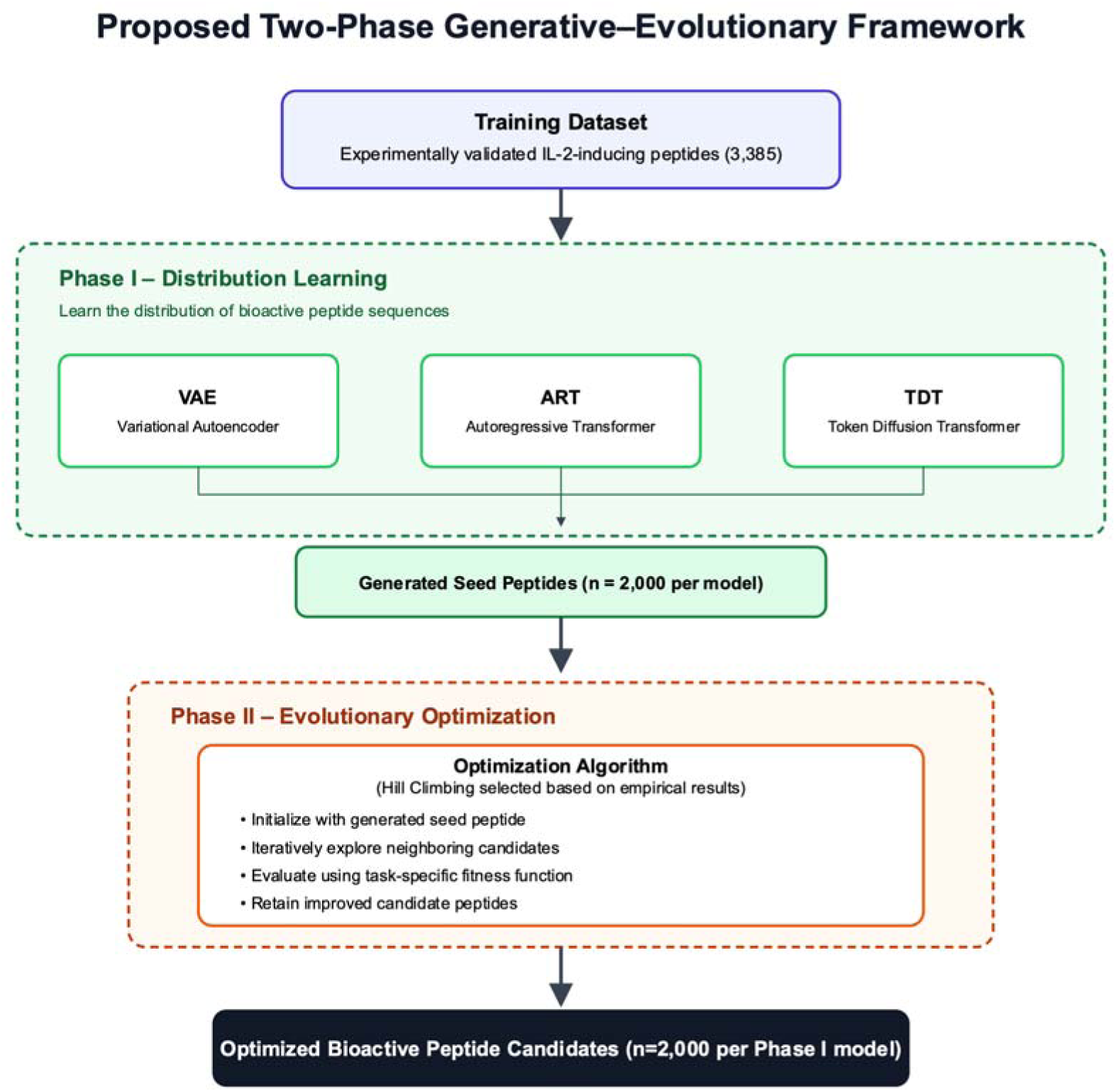
Overview of the proposed two-phase framework for bioactive peptide generation. In Phase I, distribution-learning models, including Variational Autoencoder (VAE), Autoregressive Transformer (ART), and Token Diffusion Transformer (TDT), generate diverse seed peptides from experimentally validated sequences. In Phase II, the seed peptides are refined using an evolutionary optimization algorithm to generate optimized bioactive peptide candidates with improved predicted activity and distributional similarity.

#### 2.5.1 Phase I: Distribution Learning

In the first phase, a distribution-learning model is trained on experimentally validated bioactive peptides to learn their sequence distribution. The trained model is then used to generate an initial set of peptide sequences, which serve as seed candidates for the optimization phase. These generated peptides retain the sequence characteristics of the training data while providing diverse starting points for further refinement. In this study, each distribution-learning model generated 2,000 seed peptide sequences for downstream optimization.

#### 2.5.2 Phase II: Evolutionary Optimization

In the second phase, the generated seed peptides are refined using an optimization algorithm. Each seed peptide serves as the starting point for an independent optimization process, in which candidate sequences are iteratively modified using algorithm-specific search operations and evaluated with a task-specific fitness function. The candidates are updated according to the optimization strategy to progressively improve the desired biological property while retaining information from the distribution-learning phase. Although the proposed framework is compatible with a wide range of optimization algorithms, Hill Climbing was used in the present study because it showed the best empirical performance in the comparative evaluation. Consequently, each seed peptide was refined using Hill Climbing for a maximum of 200 optimization iterations, producing the final set of 2,000 optimized peptide candidates for each Phase I model employed.

### 2.6 Evaluation Measures

#### 2.6.1 IL-2 induction confidence score

During sequence generation, optimization was guided by the IL2PepScan [9] prediction model based on the DDE feature set. Since using the same predictor for evaluation can bias the assessment, two in-silico evaluation models were used to assess the IL-2 inducing potential of the generated peptides. First, all peptides were evaluated using a separately trained Extra Trees classifier built on DPC feature set using the IL2PepScan training dataset. This is referred to as IL-2-PEM (IL-2 Primary Evaluation Model). For each generative method, the mean confidence score of the top 200 highest-scoring peptides (“Top200 Mean”) was calculated. To further assess the generalizability of the generated peptides, the same metric was also computed using the IL2Pred webserver [8], an independently developed IL-2 prediction tool. Using two predictors help determine whether the generated peptides perform well across different models rather than being optimized for a single classifier.

#### 2.6.2 KL Divergence

To evaluate whether the generated peptides reproduce the structure of real peptides beyond simple sequence similarity, we compared the distributions of compositional and physicochemical features between real and generated peptide sets using Kullback-Leibler (KL) divergence [28]. Because peptide-derived features can be high dimensional and correlated, we approximated their joint distribution using multivariate Gaussian model in a reduced PCA feature space [29]. Given two Gaussian distributions 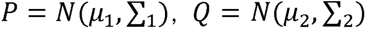 the KL divergence between them is defined as:

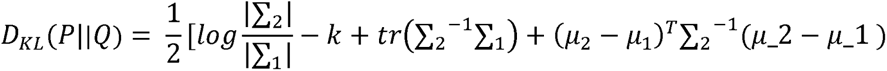

Where k denotes the dimensionality of feature space.

Since the KL divergence is asymmetric, we computed the symmetric KL divergence –

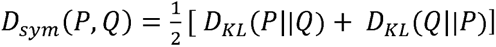

Lower symmetric KL values indicate closer agreement between the statistical distributions of generated and real peptides.

#### 2.6.3 Novelty

To quantify whether the generated peptides are different from known IL-2 inducing peptides. we computed sequence novelty using normalized Levenshtein edit distance, a widely used metric for comparing biological sequences [30]. For two sequences s1 and s2, the normalized edit distance was defined as –

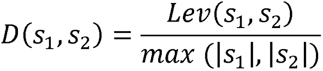

Where Lev(*s_i_*, *s_j_*) denotes the Levenshtein distance and |*s_i_*| denotes the sequence length.

For each generated peptide *g_i_*,

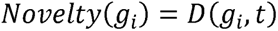

And the overall novelty score is,

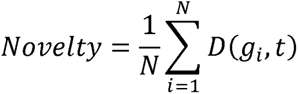

where T denotes the training peptide set and N is the number of generated peptides.

Higher novelty values indicate that generated peptides differ significantly from known training sequences, suggesting exploration of new regions of sequence space rather than memorization.

#### 2.6.4 Diversity

To assess the redundancy within the generated set, we computed intra-set diversity as the mean pairwise normalized edit distance between generated sequences. Given a set of generated peptides G, diversity was defined as –

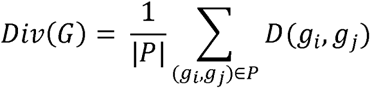

Where P is the set of sampled sequence pairs and D (*g_i_*, *g_j_*), is normalized edit distance defined above. For computational efficiency, pairwise distances were estimated using random sampling of sequence pairs up to a fixed upper limit. Higher diversity values indicate that the generator does not collapse to a limited set of sequences and instead produces structurally varied candidates.

## 3. RESULTS

### 3.1 Results of Standalone Methods

#### 3.1.1 Predictive performance

The predictive performance of peptides generated by standalone distribution-learning and optimization-based methods is summarized in the Table 1. The generated sequences were evaluated using two predictors i.e IL-2-PEM and IL2Pred, as discussed above. As shown in Table 1, Hill Climbing consistently outperformed all other methods across the two prediction models and the average predictive performance is reported for top 10% of the generated peptides (n=200). Among distribution-learning models, the Autoregressive Transformer demonstrated the strongest performance. Its performance (0.79 on IL-2-PEM and 0.88 on IL2Pred) indicates its ability to generate high-quality candidates, albeit less consistently than optimization-based approaches. Other methods, including the Variational Autoencoder and Token Diffusion Transformer showed relatively modest performance.

**Table 1.** Comparison of mean confidence scores for the top 200 peptides generated by standalone Hill Climbing (HC), Genetic Algorithm (GA), Variational Autoencoder (VAE), Autoregressive Transformer (ART), and Token Diffusion Transformer (TDT) using the IL-2 Primary Evaluation Model (IL-2-PEM) and IL2Pred (Best scores are shown in bold).

| Algorithm | IL-2-PEM Mean Score<br>(Top 200) | IL2Pred Mean Score<br>(Top 200) |
| --- | --- | --- |
| **HC** | **0.84** | **0.93** |
| **GA** | 0.69 | 0.84 |
| **VAE** | 0.74 | 0.84 |
| **ART** | 0.79 | 0.88 |
| **TDT** | 0.75 | 0.84 |

It can be observed that Hill Climbing was the best-performing optimization-based method, achieving a mean score of 0.84 on IL-2-PEM and 0.93 on IL2Pred. This suggests that hill climbing efficiently exploits the learned fitness landscape to preferentially sample high-scoring regions. In comparison, the Genetic Algorithm showed comparatively poorer performance, achieving mean scores of 0.69 on IL-2-PEM and 0.84 on IL2Pred, both lower than those achieved by Hill Climbing.

Notably, the ranking of methods was largely preserved across both IL2PepScan and IL2Pred evaluations. This consistency suggests that the observed improvements are not specific to a single predictive model. In particular, the strong performance of Hill Climbing on the DPC-based IL-2-PEM prediction model, despite being optimized using DDE features, indicates that the optimization process captures signals that generalize across feature representations.

#### 3.1.2 Distributional similarity to experimentally validated peptides

In order to evaluate how closely generated peptides resemble real IL-2–inducing peptides that have been tested experimentally, we compared feature distributions using KL divergence in physicochemical (PCP) space. Because peptide activity is largely determined by physicochemical properties rather than exact amino acid sequences, distributional similarity in PCP space serves as a meaningful indicator of biological realism. As shown in table 2, the Token Diffusion Transformer achieved the lowest KL divergence (0.10), indicating the closest match to real peptide distributions. The Variational Autoencoder and Autoregressive Transformer also showed low divergence values, indicating that they effectively captured the underlying sequence distribution. In contrast, the optimization-based methods exhibited substantially higher divergence. Hill Climbing produced a KL divergence of 2.26, while the Genetic Algorithm showed even larger deviations. These results suggest that although optimization-based methods generate high-scoring peptides, they tend to move away from the distribution of experimentally validated bioactive peptides. A similar trend is observed in Figure 2, where the physicochemical property distributions of peptides generated by the distribution-learning models closely match those of the real peptides, whereas the optimization-based methods show noticeable shifts. These findings further support that distribution-learning models are more effective at preserving overall sequence characteristics.

**Table 2.**
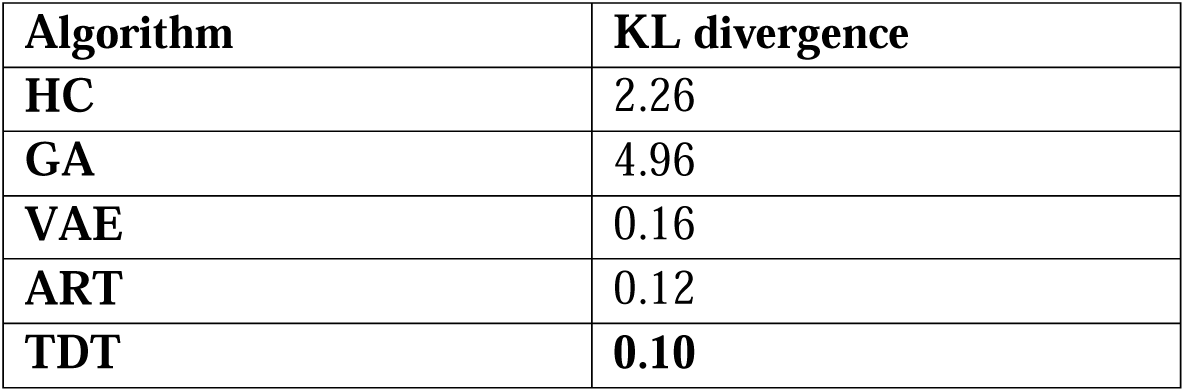
Comparison of peptide distribution similarity for standalone Hill Climbing (HC), Genetic Algorithm (GA), Variational Autoencoder (VAE), Autoregressive Transformer (ART), and Token Diffusion Transformer (TDT), measured using Kullback–Leibler (KL) divergence. Lower values indicate greater similarity to the training peptide distribution. Best scores are shown in bold.

**Figure 2.**
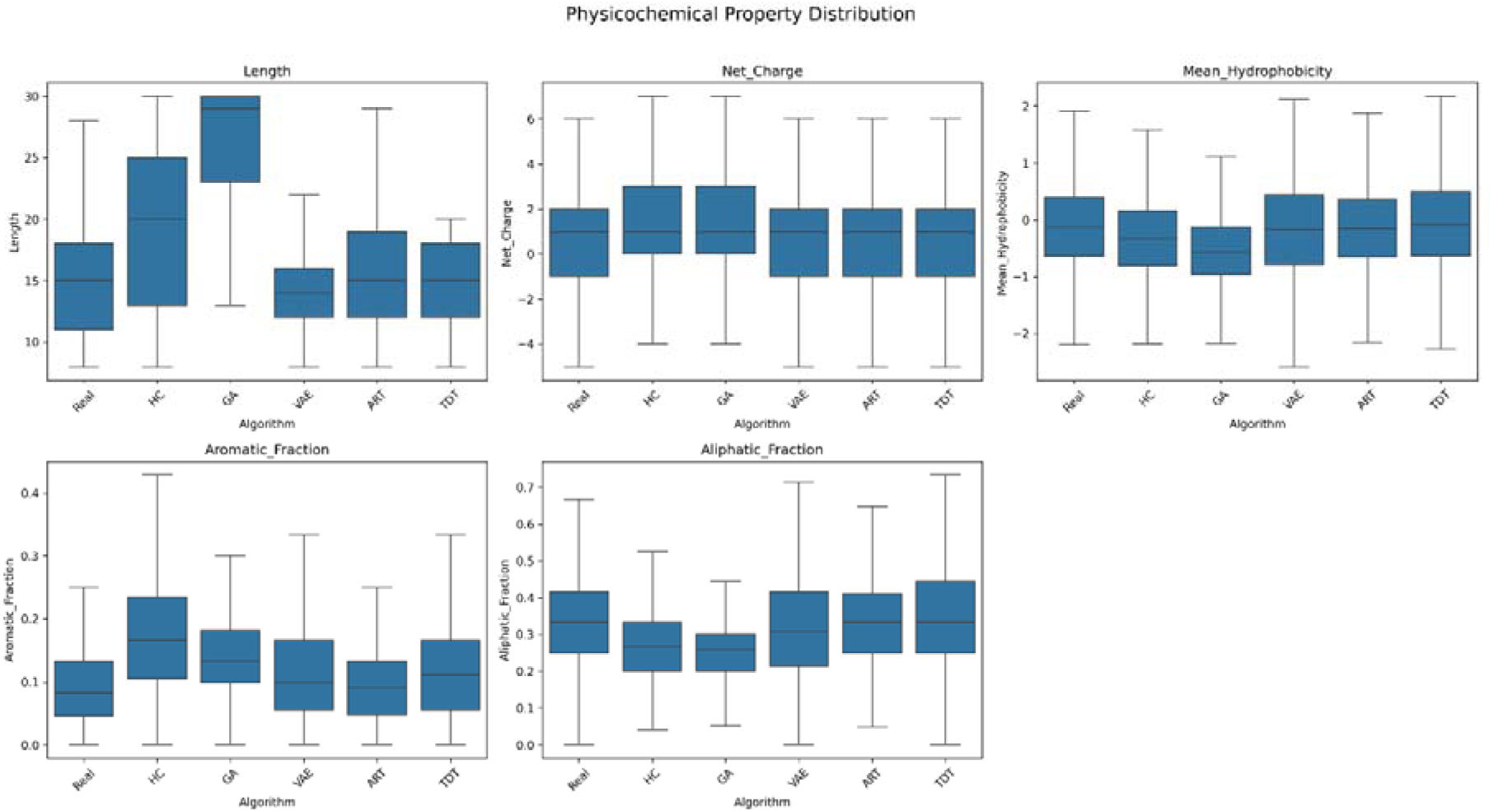
Comparison of the physicochemical property distributions of experimentally validated IL-2–inducing peptides and peptides generated by Hill Climbing (HC), Genetic Algorithm (GA), Variational Autoencoder (VAE), Autoregressive Transformer (ART), and Token Diffusion Transformer (TDT). Box plots show the distributions of peptide length, net charge, mean hydrophobicity, aromatic residue fraction, and aliphatic residue fraction. The x-axis represents the peptide source, and the y-axis represents the corresponding physicochemical property value. These properties were used to assess how well the generated peptides preserve the characteristics of the training peptide distribution.

#### 3.1.3 Novelty and Diversity of Generated Peptides

The novelty and diversity of the generated peptides provide insight into how extensively each method explores the sequence space. The Genetic Algorithm achieved the highest novelty score (0.67), followed by Hill Climbing (0.63), indicating that these methods generated sequences that were more distinct from the training peptides. In contrast, the Variational Autoencoder and Transformer-based models showed slightly lower novelty values (0.57–0.59), suggesting that they remained closer to the learned sequence distribution (Table 3). Despite these differences, all methods maintained high diversity, with values ranging from 0.85 to 0.87. The Variational Autoencoder achieved the highest diversity (0.87), which was slightly higher than that of the training dataset (0.86). This suggests that distribution-learning models can generate diverse peptide sequences without repeatedly producing similar sequences. Overall, these results suggest a clear distinction that optimization-based methods favor exploration of novel regions, while distribution-learning approaches emphasize fidelity to the underlying data distribution, all while maintaining comparable levels of diversity.

**Table 3.** Comparison of novelty and diversity data for peptides generated by standalone Hill Climbing (HC), Genetic Algorithm (GA), Variational Autoencoder (VAE), Autoregressive Transformer (ART), and Token Diffusion Transformer (TDT). Novelty is measured as the average normalized Levenshtein distance between generated peptides and the closest peptide in the training dataset, while diversity is measured as the average pairwise normalized Levenshtein distance among generated peptides. Higher values indicate greater novelty and diversity. Best scores are shown in bold.

| Algorithm | Novelty | Diversity |
| --- | --- | --- |
| HC | 0.63 | 0.86 |
| GA | **0.67** | 0.85 |
| VAE | 0.59 | **0.87** |
| ART | 0.58 | 0.86 |
| TDT | 0.59 | 0.86 |

#### 3.1.4 Comparison of Distribution-Learning and Optimization-Based Methods

The preceding analyses demonstrate that optimization-based and distribution-learning methods exhibit distinct performance characteristics. Distribution-learning models, especially transformer-based architectures, closely reproduce the statistical properties of real peptides but achieve comparatively lower functional scores. In contrast, optimization-based methods, particularly Hill Climbing, achieve high prediction scores and generate a large number of high-confidence peptides, but fail to capture the true distribution of IL-2 inducing peptides. This distinction is also evident in Figure 3, where the optimization-based and distribution-learning methods occupy distinct regions of the performance space. These findings suggest that maximizing classifier scores alone does not produce biologically realistic peptide sequences, while distribution learning alone does not necessarily generate peptides with optimal functional properties. of the Kullback–Leibler (KL) divergence between the generated and training peptide distributions.

**Figure 3.**
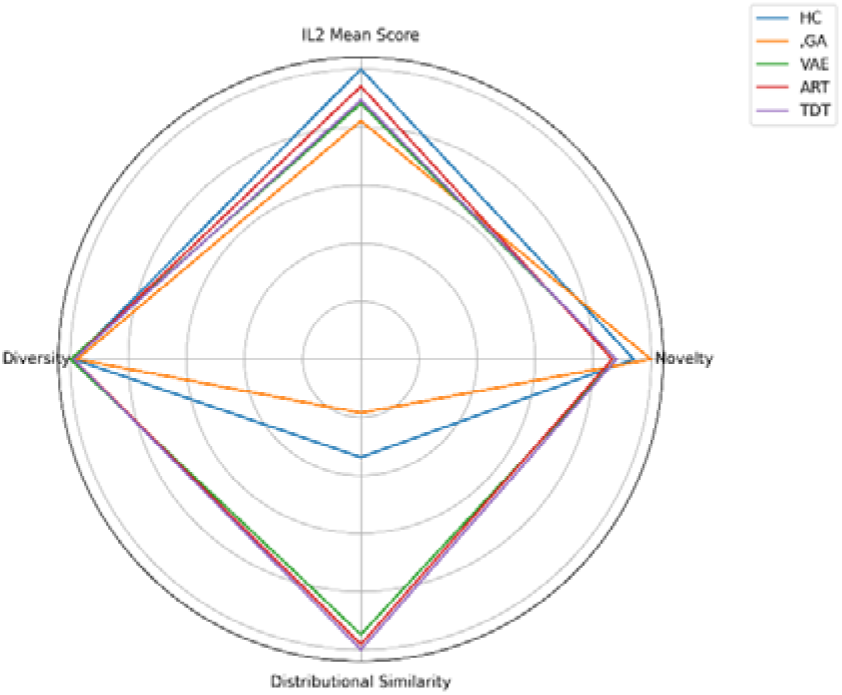
Multi-objective comparison of Hill Climbing (HC), Genetic Algorithm (GA), Variational Autoencoder (VAE), Autoregressive Transformer (ART), and Token Diffusion Transformer (TDT) across four evaluation metrics: predicted IL-2 induction score (IL-2-PEM), novelty, diversity, and distributional similarity. The radar plot summarizes the relative performance of each method, with values farther from the center indicating better performance for the corresponding metric. Novelty and diversity were calculated using normalized Levenshtein distance, while distributional similarity was calculated as the inverse

### 3.2 Results of the Proposed Two-Phase Approach: Integrating Generative Models with Optimization

To address the limitations of standalone approaches, two-phase approaches were investigated by initializing Hill Climbing with sequences generated by distribution-learning models. The goal of this approach was to combine the distributional characteristics learned by generative models with the optimization capability of evolutionary search. When initialized with sequences from the Autoregressive Transformer (Table 4), Hill Climbing achieved a mean score 0.89 (top200) on IL-2-PEM, along with 0.96 (top200) on IL2Pred. In addition to improved prediction scores, this configuration also showed substantially lower KL divergence (0.75) compared to standalone Hill Climbing (Table 2). A similar pattern was observed for initialization using the Token Diffusion Transformer (Table 4), which achieved a mean IL2Pred score of 0.95, while maintaining improved distributional alignment (KL = 0.95). Initialization with the Variational Autoencoder (Table 4) also led to improved prediction performance compared to standalone generative models.

**Table 4.** Performance evaluation of the proposed two-phase peptide generation framework, where Hill Climbing (HC) is initialized with peptides generated by Variational Autoencoder (VAE), Autoregressive Transformer (ART), and Token Diffusion Transformer (TDT). The top 200 generated peptides are evaluated using the IL-2-PEM, IL2Pred, novelty, diversity, and KL divergence. Best scores are shown in bold.

| Algorithm | IL-2-PEM Mean Score (Top200) | IL2Pred Mean Score (Top200) | Novelty | Diversity | KL Divergence |
| --- | --- | --- | --- | --- | --- |
| VAE + HC | 0.85 | 0.94 | **0.59** | **0.88** | 0.9 |
| **ART + HC** | **0.89** | **0.96** | **0.59** | 0.87 | **0.75** |
| **TDT + HC** | 0.87 | 0.95 | **0.59** | 0.87 | 0.95 |

Across all two-phase configurations, diversity remained consistently high (approximately 0.87–0.88, Table 4), comparable to standalone methods (Table 3). This indicates that improvements in prediction performance and distributional alignment were not accompanied by a loss of sequence variability. The effect of two-phase approach was also evident in the physicochemical properties shown in figure 4, where peptides generated using two-phase approaches exhibit closer agreement with real peptide properties compared to those generated by standalone optimization methods. The overall performance across evaluation metrics is illustrated in figure 5 for Hill Climbing, the Autoregressive Transformer, and their combination. These three methods were selected as representative examples of optimization-based, distribution-learning, and two-phase paradigms, respectively, to enable clearer visualization and comparison. As shown, the two-phase approach achieves a more balanced performance profile relative to the individual methods.

**Figure 4.**
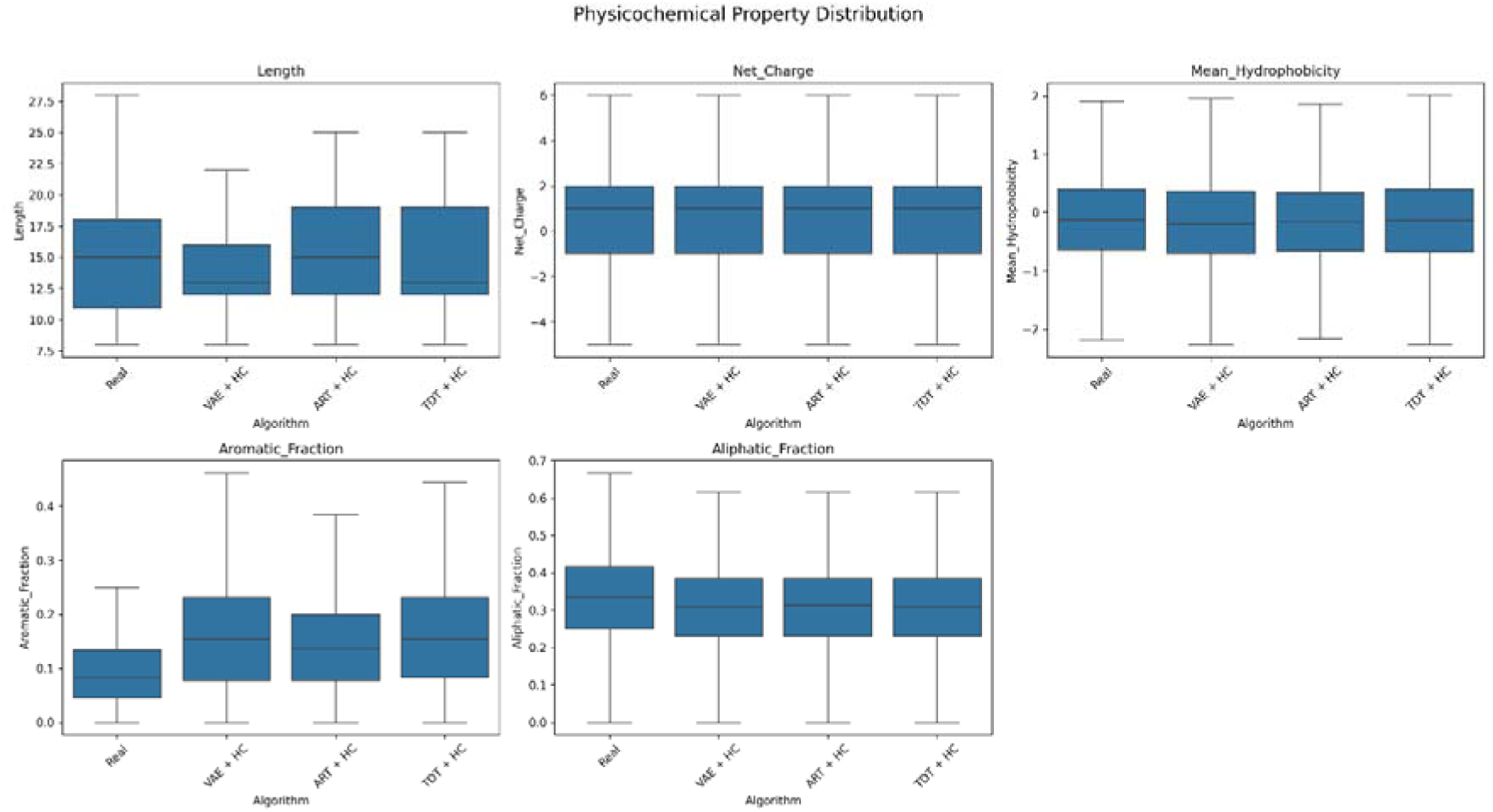
Comparison of the physicochemical property distributions of experimentally validated IL-2–inducing peptides and peptides generated by the proposed two-phase framework: Variational Autoencoder-initialized Hill Climbing (VAE + HC), Autoregressive Transformer-initialized Hill Climbing (ART + HC), and Token Diffusion Transformer-initialized Hill Climbing (TDT + HC). Box plots show the distributions of peptide length, net charge, mean hydrophobicity, aromatic residue fraction, and aliphatic residue fraction. The x-axis represents the peptide source, and the y-axis represents the corresponding physicochemical property value. The figure illustrates how well the two-phase framework preserves the physicochemical characteristics of the training peptide distribution after evolutionary optimization.

**Figure 5.**
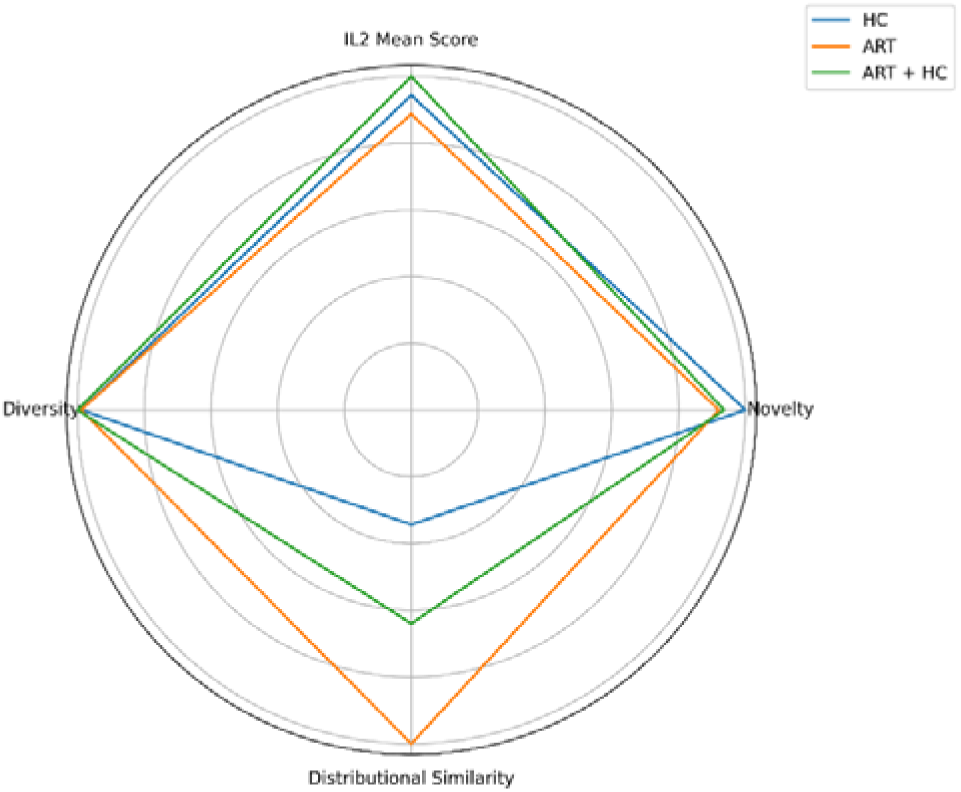
Multi-objective comparison of Hill Climbing (HC), Autoregressive Transformer (ART), and the proposed two-phase framework (ART + HC) across four evaluation metrics: predicted IL-2 induction score (IL-2-PEM), novelty, diversity, and distributional similarity. The radar plot summarizes the relative performance of each method, with values farther from the center indicating better performance for the corresponding metric. Novelty and diversity were calculated using normalized Levenshtein distance, while distributional similarity was calculated as the inverse of the Kullback–Leibler (KL) divergence between the generated and training peptide distributions.

Overall, these results demonstrate that the two-phase approach effectively combines the strengths of distribution learning and optimization for peptide generation. The proposed framework enables the generation of peptide candidates with improved functional properties while preserving the characteristics of experimentally validated bioactive peptides, making them suitable for further experimental validation.

## 4. CASE STUDY: Validation of Two-Phase Approach on an Alternative Dataset

The proposed two-phase framework generated peptides with high IL-2 induction confidence while closely matching the training distribution. Since the IL-2 dataset is relatively larger than many experimentally validated peptide datasets, the framework was further evaluated on the smaller IL-13-inducing peptide dataset to assess its generalizability under low-data conditions. This dataset was chosen owing to the availability of multiple published algorithms for the classification of these peptides [8, 9]. The IL-13 inducing peptide dataset was taken from IL13Pred [8] study which consisted of 313 IL-13 inducing peptides. All the generation methods were used to generate 2000 peptides. Optimization-based methods used the XGBoost model from iIL13Pred [9] study as the fitness function. The evaluation was done using a Random Forest model, referred to as IL-13-PEM (IL-13 Primary Evaluation Model), and the IL13Pred model to measure the IL-13 inducing potential confidence.

The evaluation scores are presented in Tables 5 and 6. For consistency with the IL-2 experiments, Hill Climbing was retained as the optimization component of the two-phase framework.

**Table 5.** Comparative evaluation of standalone Hill Climbing (HC), Genetic Algorithm (GA), Variational Autoencoder (VAE), Autoregressive Transformer (ART), and Token Diffusion Transformer (TDT) for peptide generation on the IL-13 dataset. The top 200 generated peptides are evaluated using the IL-13-PEM, IL13Pred, novelty, diversity, and KL divergence. Best scores are shown in bold.

| Algorithm | IL-13-PEM Mean Score (Top 200) | IL13Pred Mean Score (Top 200) | Novelty | Diversity | KL Divergence |
| --- | --- | --- | --- | --- | --- |
| HC | **0.92** | **0.99** | 0.69 | 0.86 | 1.76 |
| GA | **0.92** | 0.98 | **0.72** | 0.85 | 4.49 |
| VAE | 0.76 | 0.94 | 0.68 | **0.87** | 0.88 |
| ART | 0.79 | 0.96 | 0.66 | 0.86 | **0.24** |
| TDT | 0.75 | 0.95 | 0.65 | 0.89 | 0.3 |

**Table 6.** Performance evaluation of the proposed two-phase peptide generation framework, where Hill Climbing (HC) is initialized with IL-13 inducing peptides generated by Variational Autoencoder (VAE), Autoregressive Transformer (ART), and Token Diffusion Transformer (TDT). The top 200 generated peptides are evaluated using the IL-13-PEM, IL13Pred, novelty, diversity, and KL divergence. Best scores are shown in bold.

| Algorithm | IL-13-PEM Mean Score (Top 200) | IL13Pred Mean Score (Top 200) | Novelty | Diversity | KL Divergence |
| --- | --- | --- | --- | --- | --- |
| VAE + HC | 0.71 | 0.93 | **0.68** | 0.87 | 0.98 |
| ART + HC | **0.90** | **0.99** | 0.67 | 0.86 | 0.59 |
| TDT + HC | 0.72 | 0.98 | 0.65 | **0.88** | **0.36** |

A similar trend was observed for the IL-13 dataset. As shown in Table 5, the optimization-based methods achieved high IL-13 induction confidence (0.92 on IL-13-PEM and 0.98–0.99 on IL13Pred) but exhibited relatively high KL divergence (1.76–4.49), indicating that the generated peptides deviated from the physicochemical distribution of experimentally validated peptides. The distribution learning algorithms on the other hand show mean IL-13 inducing confidence in the range of 0.75 –0.79 on IL-13-PEM. This is significantly lower than optimization-based models but distribution learning approaches show lower KL divergences (0.24–0.88), with lowest being that of Autoregressive Transformer (KL = 0.24). Table 6 summarizes the evaluation scores of the two-phase approach, where Hill Climbing is initialized using peptides generated by distribution-learning methods. In the low-data regime, the Variational Autoencoder and Token Diffusion Transformer-based initializations produced comparatively lower-confidence peptides, although TDT-initialized Hill Climbing achieved the lowest KL divergence (0.36). However, in this case also, the Hill Climbing method initialized with Autoregressive Transformer is able to generate high confidence peptides closely matching the performance of Hill Climbing alone, while also significantly improving the KL divergence with a value of 0.59.

These results are consistent with those observed for the IL-2 dataset and demonstrate that ART initialized Hill Climbing effectively balances optimization performance and distribution preservation. Furthermore, its strong performance on both the larger IL-2 dataset and the considerably smaller IL-13 dataset suggests that the proposed two-phase approach is robust across different dataset sizes.

## 5. DISCUSSION

Although artificial intelligence has been widely applied to bioactive peptide discovery, most studies focus either on peptide prediction or on generating candidate sequences using individual computational approaches. Recent advances in distribution-learning models and evolutionary optimization have improved the exploration of the vast peptide sequence space. However, existing approaches typically focus on either distributional realism or functional optimization rather than both. Distribution-learning models such as Variational Autoencoders and Transformer-based architectures reproduce the statistical properties of known peptides and generate biologically realistic sequences. However, they do not consistently generate peptides with high predicted biological activity. In contrast, optimization-based methods, particularly Hill Climbing, maximize prediction scores and produce a large number of high-confidence peptides. However, these methods show poor agreement with the distribution of experimentally validated peptides, indicating that they explore regions of the sequence space that may not be biologically realistic. These observations suggest that distribution-learning and optimization-based methods have complementary strengths and motivate their integration into a single framework.

This study systematically compared multiple generative approaches for IL-2–inducing peptide generation and identified a common limitation of standalone methods. Optimization was performed using the IL2PepScan model, whereas evaluation was carried out using a separately trained IL-2-PEM model [9] together with an independent predictor, IL2Pred [8]. The consistent trends across both evaluation models indicate that the observed improvements are not restricted to a single predictor. Instead, they suggest meaningful enrichment of IL-2–inducing peptides. The ability of the optimization-based methods to perform well across different feature representations further indicates that the learned fitness landscape captures general properties of bioactive peptides rather than model-specific patterns. On the IL-2 dataset, ART-initialized Hill Climbing retained high predictive performance (IL2Pred = 0.96, IL-2-PEM = 0.89) while reducing the KL divergence from 2.26 to 0.75, demonstrating that high-confidence peptide generation can be achieved without compromising distributional similarity.

The IL-13 case study further supports these findings. Although the dataset contained only 313 experimentally validated peptides, the proposed framework maintained high predictive performance while preserving distributional similarity. ART-initialized Hill Climbing achieved IL13Pred and IL-13-PEM scores of 0.99 and 0.90, respectively, while reducing the KL divergence from 1.76 to 0.59. These results indicate that the framework performs well even when the amount of training data is limited and may be useful for peptide discovery problems where experimentally validated peptides are scarce.

While recent comparative studies [6, 31] have evaluated different peptide generation paradigms independently, they do not explicitly integrate distribution-learning and optimization-based methods within a unified framework. The proposed two-phase framework addresses this limitation by using distribution-learning models to generate seed peptides, followed by optimization-based refinement. As a result, the generated peptides show improved prediction scores while remaining consistent with the distribution of experimentally validated peptides. The improvements are not limited to prediction performance but are also reflected in distributional similarity and sequence diversity.

These findings indicate that the choice of initialization affects optimization performance. Random initialization may guide the search towards unrealistic regions of the sequence space, whereas initialization using a distribution-learning model provides a more suitable starting point. Although demonstrated using Hill Climbing, this observation is not specific to the optimization method used in this study and may also apply to other optimization algorithms and biological sequence design problems. These observations can be explained by the exploration–exploitation trade-off [32], where distribution-learning models explore the sequence space, while optimization-based methods refine generated peptides.

The proposed framework may also be useful for addressing class imbalance in biological datasets. Unlike conventional data augmentation methods, which mainly increase the number of minority-class samples, the proposed framework generates synthetic peptides that preserve the distributional characteristics of experimentally validated bioactive peptides while exhibiting high predicted biological activity. Consequently, the generated peptides can be used both as candidate sequences for experimental validation and as biologically meaningful synthetic data for training prediction models.

Despite these encouraging results, several limitations remain. First, the generated peptides were evaluated only using computational prediction models, which cannot fully capture the complexity of biological systems. Although multiple prediction models were used to reduce dependence on a single predictor, experimental validation is required to confirm the biological activity of the generated peptides. Second, the current framework optimizes only a single property, namely IL-2 induction. Although it also performed well on the IL-13 dataset, it does not consider other important properties such as peptide stability, toxicity, or structural characteristics that influence therapeutic potential. Future work should extend the framework to multi-objective optimization by considering several biological properties simultaneously. Incorporating structural modelling and experimental feedback would further improve the practical applicability of the framework.

In conclusion, this study demonstrates that integrating distribution learning with evolutionary optimization provides an effective solution to the challenges of peptide generation and dataset imbalance. By combining the strengths of distribution learning and functional optimization, the proposed two-phase framework enables the generation of peptides that are both realistic and high-performing. This approach is generalizable and can be extended to other classes of bioactive peptides, offering a scalable and biologically informed strategy for next-generation peptide discovery.

## Code Availability

The code is available in zenodo https://zenodo.org/records/21069840

## Author Contribution

**Rachit Abhigyan-** Conceptualization, Methodology, Software, Investigation, Writing Original Draft. **Vikas Sood-** Supervision, Project Administration, Writing Review and Editing. **Pooja Arora-** Supervision, Project Administration, Funding Acquisition Writing Review and Editing and **Baljeet Kaur-** Supervision, Project Administration, Writing Review and Editing

## Use of AI

The authors acknowledge the use of artificial intelligence (AI)-based tools to assist in language refinement, grammar correction, and overall readability of the manuscript. The AI tools were used solely to improve clarity and presentation only. All scientific ideas, analyses, and interpretations presented in this work are the original contributions of the authors.

## Ethics statement

All the data used in this work is publicly available and no new data was generated during this work.

